# From Context to Code: Rational De Novo DNA Design and Predicting Cross-Species DNA Functionality Using Deep Learning Transformer Models

**DOI:** 10.1101/2023.10.15.562386

**Authors:** Gurvinder Singh Dahiya, Thea Isabel Bakken, Maxime Fages-Lartaud, Rahmi Lale

**Affiliations:** Department of Biotechnology and Food Science, Faculty of Natural Sciences, Norwegian University of Science and Technology, Trondheim, Norway; Syngens AS, Trondheim, Norway

**Keywords:** Artificial Intelligence, Synthetic Biology, Bacteria, Coding sequence, Promoter, 5′ untranslated region

## Abstract

Synthetic biology currently operates under a framework dominated by trial-and-error approaches, which hinders the effective engineering of organisms and the expansion of large-scale biomanufacturing. Motivated by the success of computational designs in areas like architecture and aeronautics, we aspire to transition to a more efficient and predictive methodology in synthetic biology. In this study, we report a DNA Design Platform that relies on the predictive power of Transformer-based deep learning architectures. The platform transforms the conventional paradigms in synthetic biology by enabling the context-sensitive and host-specific engineering of 5′ regulatory elements—promoters and 5′ untranslated regions (UTRs) along with an array of codon-optimised coding sequence (CDS) variants. This allows us to generate context-sensitive 5′ regulatory sequences and CDSs, achieving an unparalleled level of specificity and adaptability in different target hosts. With context-aware design, we significantly broaden the range of possible gene expression profiles and phenotypic outcomes, substantially reducing the need for laborious high-throughput screening efforts. Our context-aware, AI-driven design strategy marks a significant advancement in synthetic biology, offering a scalable and refined approach for gene expression optimisation across a diverse range of expression hosts. In summary, this study represents a substantial leap forward in the field, utilising deep learning models to transform the conventional design, build, test, learn-cycle into a more efficient and predictive framework.

## Introduction

Achieving predictive power represents a pivotal step in synthetic biology, bridging the long-standing divide between genotype and phenotype^1^. It has the potential to move the field from being trial-and-error heavy to data-driven, informed decision-making^2^. Drawing inspiration from disciplines like architecture and aeronautics, where initial computational designs often lead to functional outcomes, we envision a similar seamlessness in synthetic biology^3^. As architects and aviation engineers craft structures and planes that work on the first attempt, recent efforts prove that the linear progression from Design to Function can alter the current framework of design, build, test, and learn-cycle. This progression sidesteps the intricate and exhaustive Design and Test phases that have long been inherent to the process^4^.

While computational approaches are being developed, engineering living systems using synthetic biology methods still heavily relies on trial-and-error approaches mostly limited to single model organisms. The assembly of individual—pre-characterised—DNA parts, akin to a LEGO construction, often neglects the crucial aspect of host- and composition-specific DNA context^5^, leading to sub-optimal outcomes in gene expression, protein production, and overall system performance. Consequently, laborious high-throughput screening efforts have become the brute-force method of choice, becoming impractical and cost-prohibitive for large-scale biomanufacturing needs.

To address the aforementioned challenges, we propose an approach that harnesses the predictive power of artificial intelligence. Specifically, we leverage Transformer-based architectures^6^, which are a subset of generative AI^7^, for the rational de novo design of 5′ regulatory sequences and prediction of cross-species DNA functionality in model and non-model bacterial species.

Deep learning transformer models have demonstrated exceptional capabilities in natural language processing tasks, thanks to their ability to capture intricate contextual relationships in data. These models, initially developed for processing human language data, have shown great potential for biotechnological applications beyond text analysis among others for genomic sequence analysis^8^, gene expression^9^, ‘omics analysis^10^, promoter design^11,12^.

One another key features of transformer models is the attention mechanism, which allows the model to focus on specific parts of the input that are relevant to the task at hand. This ability to selectively attend to relevant information is crucial in understanding the context in which biological systems operate, identifying key factors that may influence performance. Transformer models can also be used in generating interpretable representations of biological data, aiding researchers to understand the underlying patterns and relationships. This interpretability can aid in rationalising the design and assembly of DNA parts, considering the context-dependency. And finally, the flexibility of transformer models allows them to be adapted to different organisms and biological systems. This adaptability ensures that the models can capture the unique context of different species, considering the variations that may exist across different biological host context.

In the realm of synthetic biology, “context” refers to the intricate and often unpredictable circumstances in which biological systems operate. It encompasses a wide range of factors that can influence the behaviour and function of biological components, leading to potential discrepancies between design and actual performance. The source of contextual dependencies are defined as compositional, host, and environmental^5^. Compositional context deals with the interaction between biological elements and the challenges that emerge from their physical and functional interaction. Host context pertains to the inherent reliance of a biological system on the host organism’s biochemical capacity for its functioning. This leads to a universal linkage and rivalry between both heterologous and native functions. Environmental context stems from external variables that impact the functioning of the biological system. Understanding and controlling context is therefore a critical aspect of successful synthetic biology design and implementation, however, this aspect has proven to be a challenge in the field.

In this study, we report a DNA Design Platform that transforms the conventional paradigms in synthetic biology by enabling the context-sensitive and host-specific engineering of 5′ regulatory elements—promoters and 5′ untranslated regions (UTRs). A unique feature of the platform is its ability to handle not just a single codon optimised coding sequence (CDS) variant, which has been the traditional approach in the field, but an extensive array of CDS variants for generating the 5′ regulatory sequences and predicting the transcription and translation rates from the resulting DNA constructs. This novel approach enables us to generate unique 5′ regulatory sequences that are customised for each codon-optimised CDS variant in the target host, thereby achieving an unparalleled level of specificity and adaptability. Consequently, we substantially broaden the spectrum of possible gene expression profiles and phenotypic outcomes, providing a more refined and scalable strategy for the tailoring of genetic constructs beyond just model organisms.

The emerging paradigm of predictable DNA design offers a groundbreaking avenue to surmount many of the field’s challenges, providing the means to directly translate genetic sequences into predictable phenotypic outcomes. This development has significant implications for advancing our capabilities in fine-tuning gene expression and optimising metabolic pathways across a wide range of expression hosts. The context-aware AI-driven design approach significantly reduces reliance on trial-and-error and minimises the need for extensive high-throughput screening efforts, marking a significant advancement in the field of synthetic biology.

## Results

For the development of a rational DNA design for 5′ regulatory sequences and codon-optimised CDS variants that leads to predictable output, we have targeted the following objectives. The DNA design should:

- not be limited by solely relying on the existence of pre-characterised DNA parts;
- not be limited to a single but should consider a large set of codon-optimised coding sequences;
- overcome the reliance on model organisms and should enable the utilisation of non-model organisms;
- enhance predictability by enabling the prediction of biological outcomes to create a genotype-phenotype relationship;
- limit the need for extensive high-throughput screening and enable more targeted and streamlined experimental efforts.

Aiming to achieve these objectives, our preliminary focus has centred on the predictable design of 5′ regulatory sequences, sensitive to both host and DNA contexts, specific to codon-optimised CDSs using transformer models^6^. We taliored Transformer models specifically for DNA sequences to capture the intricate contextual interrelationships within DNA, thereby enabling us to predict the functionality of genetic constructs across a range of bacterial species.

### Evolutionary Forces on Coding and Regulatory sequences

Evolution plays a fundamental role in shaping the genomes of organisms, leading to the development of unique biochemical repertoire and capacities in each one of them^13^. While the mechanisms of transcription and translation are universally conserved, the sequences that participate in these processes exhibit a remarkable level of diversity.

Codon usage refers to the frequency with which specific codons are used to encode a particular amino acid within a gene. Within a single organism, different genes can have distinct patterns of codon usage. This intragenomic variation is influenced by a range of factors including gene expression levels, GC content, and evolutionary age of the genes. For instance, highly expressed genes often show a strong preference for specific codons that are more efficiently translated, thereby optimising protein synthesis. Several studies also focus on the amino acid composition and the use or rare codons in the start of coding sequences^14,15^. However, the codon usage exhibits both intra-species and inter-species variability. To illustrate the variances in codon usage among different bacterial species, we conducted an analysis focusing on the initial 100 codons of highly expressed genes—those in the top 30% based on RNA-seq datasets—in *Bacillus subtilis, Corynebacterium glutamicum, Escherichia coli*, and *Streptomyces venezuelae*. While the use of rare codons is notably significant in highly expressed genes for *E. coli*, its importance diminishes for the other examined bacterial hosts (Figure S1).

Presently, the standard approach to addressing codon usage disparities in synthetic biology applications is through gene synthesis, specifically by optimising the CDSs to suit the codon preferences of the target host organism. A major drawback of this practice is the reliance on a single optimised codon variant for gene expression analyses.

While certain CDSs are highly conserved across diverse species—signifying their vital importance in essential biological processes—the regulatory sequences controlling gene expression frequently exhibit substantial variability, being subject to evolutionary pressures like mutation and natural selection. Such variability in regulatory regions enables differential gene expression profiles, consequently leading to unique phenotypic characteristics. This adaptability is crucial for the survival and diversification of species across different ecological niches^16,17^.

Although codon usage diversity is well-understood and has led to the development of optimisation tools, the variability in regulatory sequences has been comparatively less explored and lacks similar computational resources. To exemplify the differences in regulatory sequences both within and across species, we chose four orthologous clusters as reference points, utilising HGTree v2.0 for this purpose^18^. The clusters selected come from the datasets acquired from the public repositories and encompass a mix of conserved genes, as well as hosts that belong to either the same genus or multiple genera. Each cluster is characterised by a unique protein, denoted by its specific amino acid sequence, but exhibits differences in the CDS due to host-dependent codon usage. The clusters contain the following proteins identified by their UniProt IDs: A0A064BZB4, A0A0E1M7N8, A0A0J6J4S5, and A0A1C1EV68. For the examination of regulatory sequences, we collected 100 base pairs upstream of the start codon from each cluster and created a WebLogo^19^ by aligning these sequences. Although the CDS for the four genes remain conserved across various species, the regulatory sequences display noticeable variation (Supplementary Figures S2, S3, S4 and S5). This is consistent with the notion that specific regulatory elements like promoters, enhancers, and transcription factor binding sites can vary substantially among different microorganisms, underscoring the need to treat each host as unique for the DNA composition of 5′ regulatory sequences and CDSs.

### Developing a DNA Design Platform

To accommodate host- and context-specific variations in both regulatory sequences and CDSs, we sought to develop a platform for either *de novo* creation or the optimisation of existing DNA sequences, ensuring a high level of predictability.

In our design framework, two key aspects significantly diverge from current practices. First, we do not separate the promoter and 5′ UTR sequences, as their compositional elements often overlap and cannot be easily dissected^20^. For example, the initially transcribed region at the 5′-end of the 5′ UTR influences the abortive profile^21^; the transcriptional and translational characteristics are shared across the promoter and the 5′ UTR sequences^22,23^. Second, when it comes to the CDS, no single design meets all the criteria used for generating codon-optimised variants^14,15,24^. Therefore, to capture these unique features, we design the 5′ regulatory sequences as an integrated DNA element and consider not just one, but a range of codon-optimised CDS variants in combination.

The process of generating context-aware sequences begins with the identification of the CDS and the expression host, which could be either a single or multiple hosts (Figure 1). From this initial data, we perform CDS optimisation specific for the host(s), allowing for not just a single but a range of codon-optimised CDS variants. Subsequently, for each codon-optimised CDS, the system designs 5′ regulatory sequences. Upon completion, we *in silico* score and rank all the resultant DNA constructs. Given the product’s nature, a subset of DNA constructs is chosen for DNA synthesis, followed by DNA transformation and expression studies in microorganisms.

**Figure 1.**
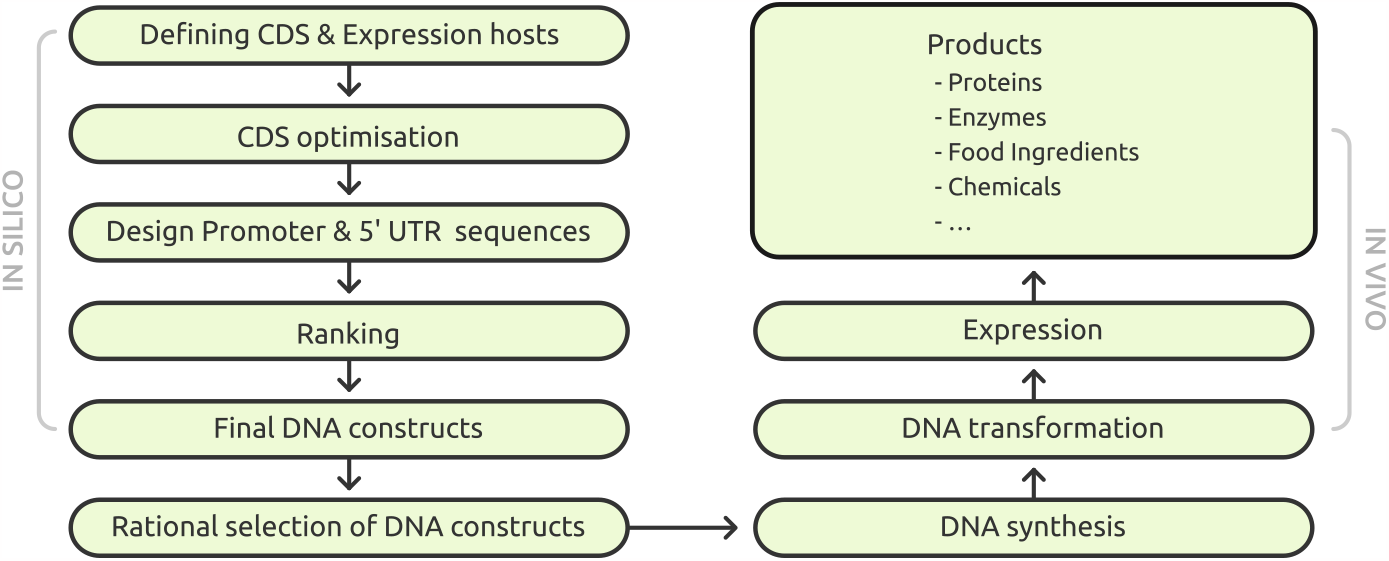
The process spans from the identification of the CDS and expression host through to the phenotypic characterisation of the DNA design and the steps of experimental validation.

**Figure 2.**
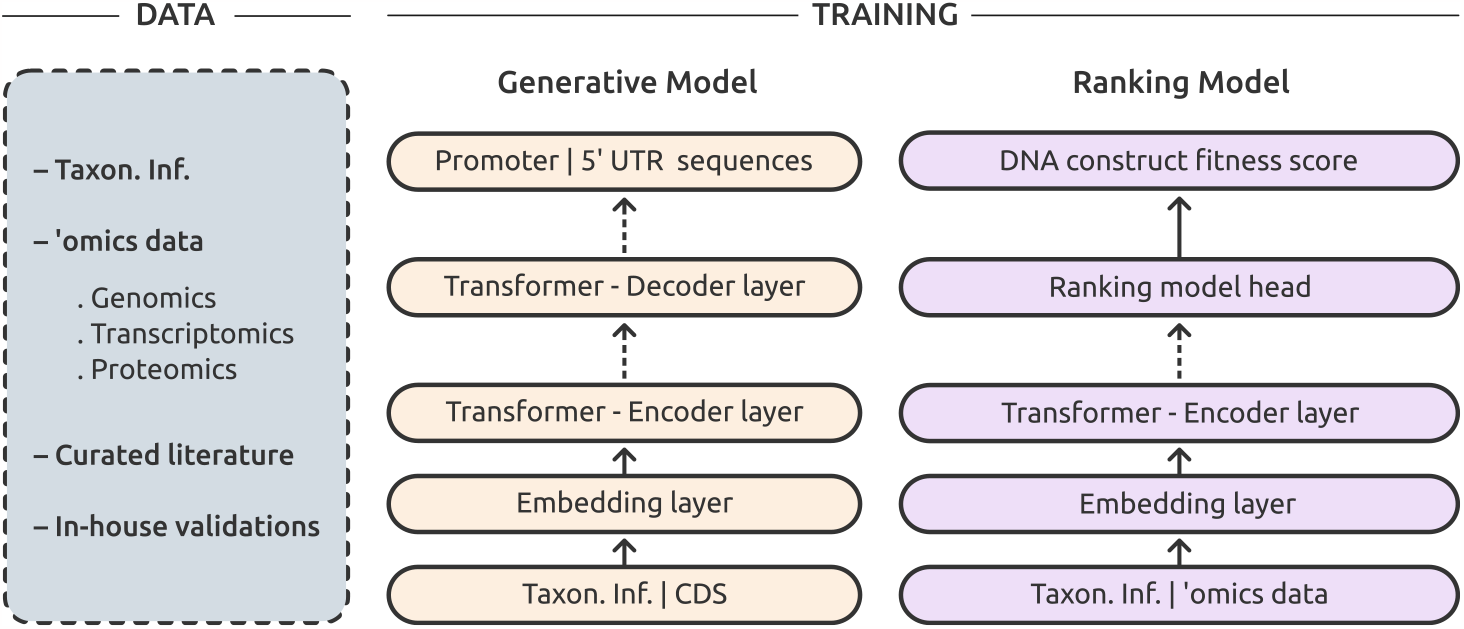
Data, training and models used.

### Data and model training

The DNA design platform integrates genomics, transcriptomics, and proteomics datasets to understand the biological characteristics of microorganisms. These input datasets are tokenised and analysed using Transformer-based deep learning models.

The platform consists of two main modules: (i) The generator model employs an encoder-decoder architecture and uses an autoregressive approach to generate 5′ regulatory sequences based on the DNA context. This context is embedded by the encoder and then relayed to the decoder layer to yield the generate 5′ regulatory sequences; (ii) The ranking model, an encoder-only construct, evaluates the provided context, which includes taxonomy information, 5′ regulatory sequences, CDS, amino acid sequences and other ‘omics data, to assess the expressibility of the specific DNA construct. This model is sensitive to even single base pair alterations and generates a fitness score for each DNA construct.

### Generation of coding sequence variants and sequence optimisation

The platform produces a multitude of optimised CDSs, drawing on taxonomic information and amino acid sequences. It offers the flexibility to optimise either the entire CDS or specific regions, based on given parameters and constraints. Following the generation of candidates, each undergoes a series of predefined checks for validation using DNA Chisel^25^. From this pool, we then evaluate and select only a few high-performing candidates based on the context provided (Figure 3).

**Figure 3.**
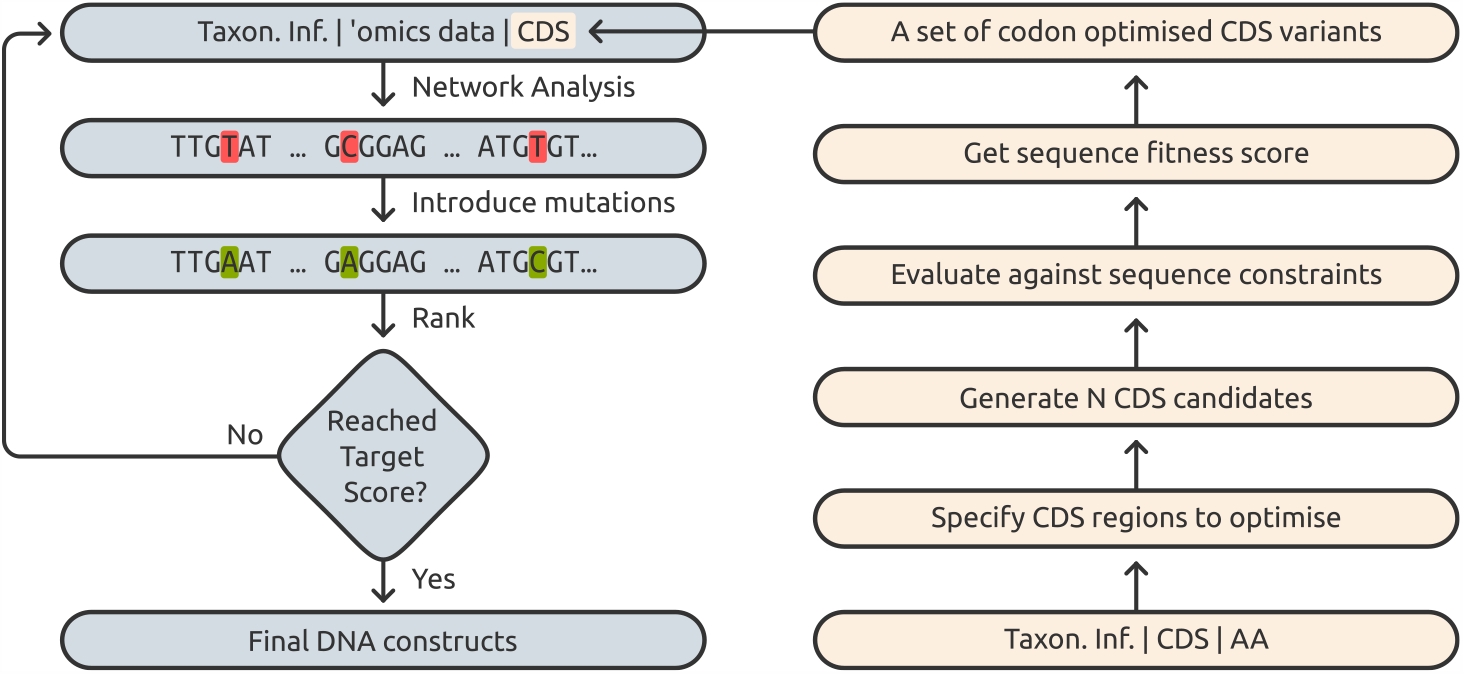
The steps involved in generating coding sequence variants and sequence optimisation.

Additionally, the platform has the capability to further refine a given DNA construct to enhance its expression. It introduces mutations at identified hotspots, based on model analysis of the existing construct, and subsequently re-scores it. This iterative process can be repeated until either the specified target is achieved or a predetermined number of cycles is reached (Figure 3).

### Constitutive expression

In the development of the platform, the first model established was the generator model. In our preliminary assessment of the generator model’s capacity, we de novo designed 24 new 5′ regulatory sequences. Specifically, we created three distinct constructs for each of four host bacterial species: *B. subtilis, C. glutamicum, E. coli*, and *P. putida*, employing two fluorescent proteins, GFP and mCherry. Of these 24 sequences, 19 were successfully constructed, all of which resulted in variable levels of fluorescent protein expression (Figure 4, Supplementary Material). Each sequence carried a Shine-Dalgarno sequence, adhering to canonical translation regulation. To assess the uniqueness of these regulatory sequences, we performed a BLAST search. The BLAST outcomes revealed no significant similarities found for any of the regulatory sequences (Figure 4). The uniqueness of the sequences is a confirmation that the generator model possess the capability to generate entirely novel sequences.

**Figure 4.**
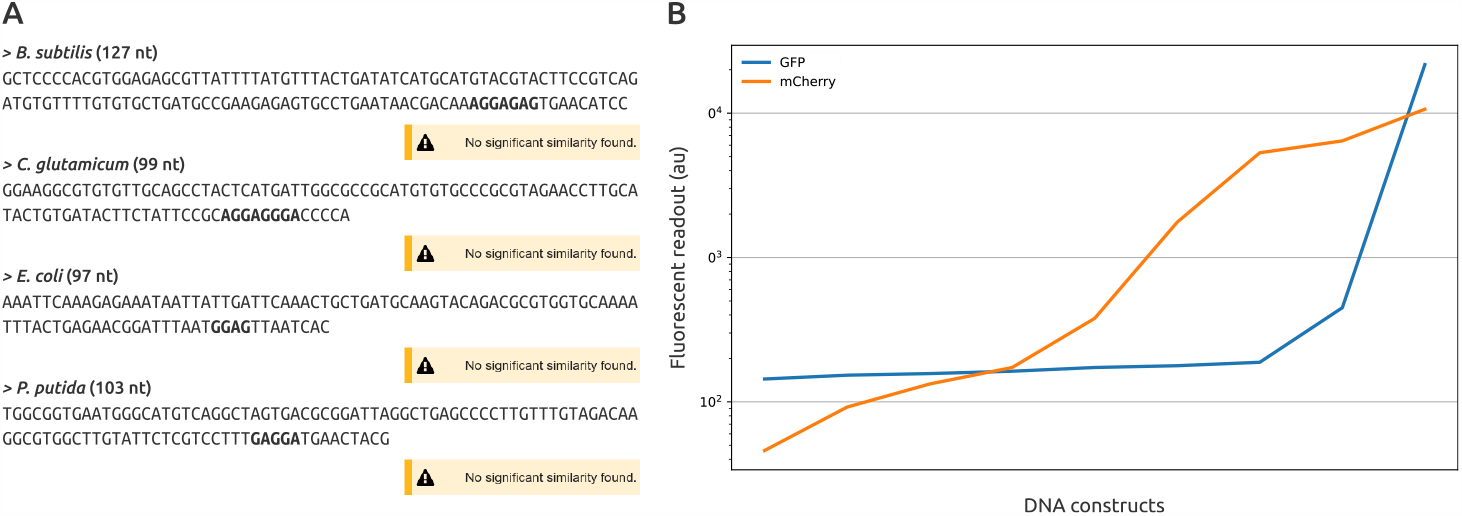
**A** Representative de novo synthesised promoter and 5′ UTR sequences with theoretical Shine-Dalgarno sequences highlighted in bold. The BLAST outcomes for each sequence are displayed below them. **B** Fluorescent protein expression profiles for 19 constructs; with three unique constructs for each host and two CDSs, examined in four bacterial species *B. subtilis, C. glutamicum, E. coli*, and *P. putida*.

### Regulatable expression

Following the successful demonstration of the generator model’s capabilities, we proceeded to develop the ranking model. To evaluate the combined effectiveness of both the generator and ranking models, we opted to fine-tune and predict the performance of an existing inducible expression system. For this task, we employed the ChnR/Pb expression system, which is positively regulated and originates from *Acinetobacter* sp.^26^. Using the native system as a base, we constructed 5′ variants that resulted in four different expression profiles, utilising GFP as the reporter gene. The generator model was promoted with the initial base subsequence and model was asked to complete the sequence. Upon generation, ranking model predicted the fitness score for each of those sequences. All sequences were evaluated and max scaled based on experimental values. Figure 5 illustrates that the developed models are capable of engineering an inducible system to achieve a spectrum of expression levels.

**Figure 5.**
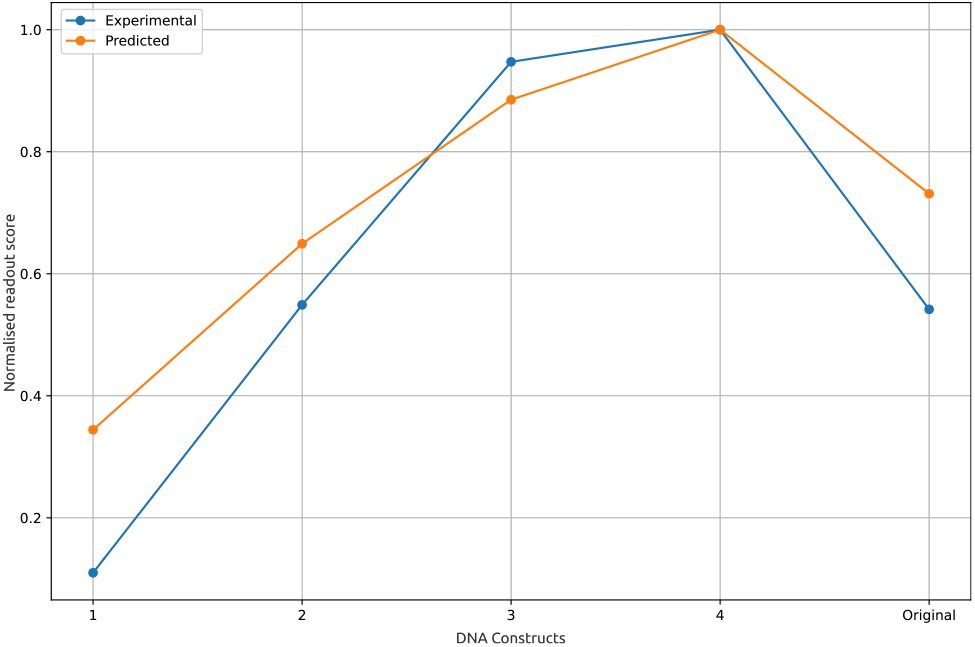
Green fluorescent protein expression profiles of four constructs compared to the original ChnR/Pb inducble expression system. The predicted levels are depicted in an orange and the experimentally verified levels are in blue colour. Spearman’s rank correlation value is 0.72 (p < 0.01) between model predictions and expression values.

### Enhanced Production of a Commercially Relevant Enzyme

As an application of the DNA design platform, we focused on Moloney Murine Leukemia Virus Reverse Transcriptase (M-MuLVRT), an industrially important enzyme. M-MuLVRT functions as an RNA-directed DNA polymerase, capable of initiating the synthesis of a complementary DNA strand from a primer, using either RNA for cDNA synthesis or single-stranded DNA as a template.

We sourced an existing construct that employs an inducible system currently utilised for the production of the enzyme in the market in *E. coli*. Given the inducible nature of the original construct, we opted to employ the ChnR/Pb system for this demonstration in *E. coli*. Initially, we used the platform to generate 105 codon-optimised variants of the M-MuLVRT CDS. Subsequently, we produced 11 705 optimised 5′ regulatory sequences for these CDS variants using the ChnR/Pb system, and computationally ranked them based on the predicted expression levels. From the highest-performing candidates, we selected five constructs and synthesised their complete gene expression cassettes in a plasmid form. Following DNA transformation and expression studies, constructs 1 and 5 emerged as the most effective (Figure S6). We then characterised both the soluble and insoluble fractions by SDS-PAGE using the DNA constructs 1 and 5 (Figure 6). Utilising semi-quantitative measurements of band intensities, we were able to confirm that the expression of M-MuLVRT was increased by 40%. This demonstratin indicated the platform’s capacity to optimise and enhance the production of industrially relevant enzymes, confirming its utility for biomanufacturing purposes.

**Figure 6.**
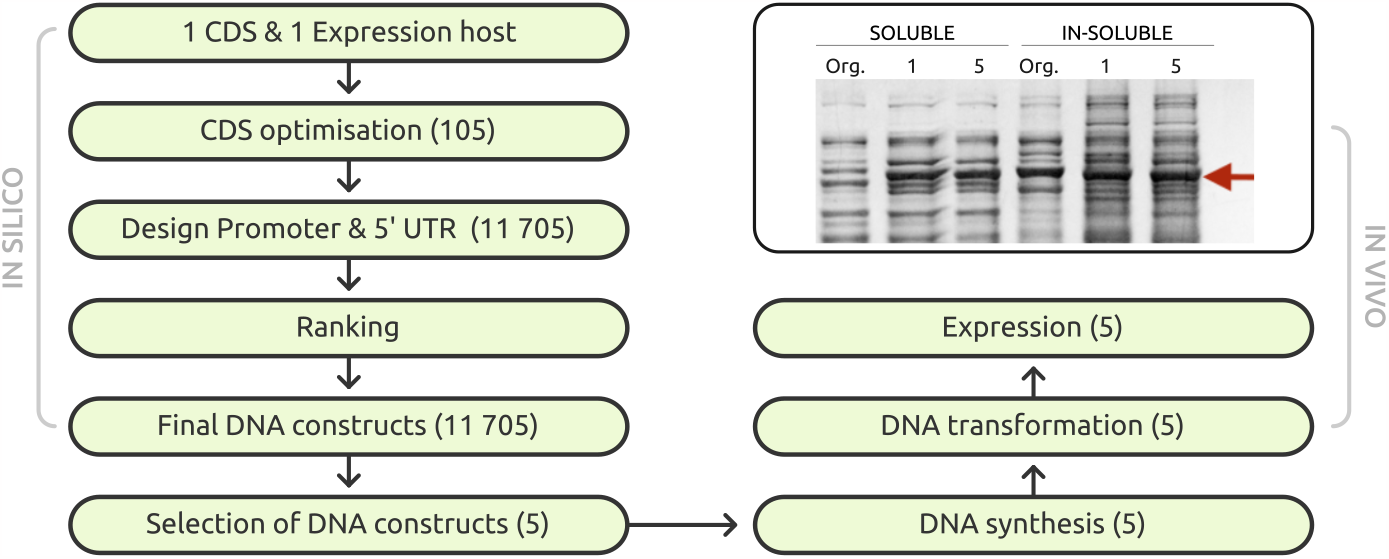
The in silico design, rational selection, DNA synthesis and characterisation of M-MulvRT expression in *E. coli* using the ChnR/Pb expression system. The image depicts the SDS-PAGE analysis of the enzyme both soluble and insoluble fractions obtained from the original (Org.) and the designed constructs 1 and 5. The red arrow indicates the expected molecular size of the enzyme, aprox. 77 kDa.

### Predicting Phenotypes from Novel Data

The results presented so far underscore the platform’s capacity in creating 5′ regulatory sequences along with CDS variants that yield predictable results. An additional feature of our platform lies in its capacity to predict the phenotype of unseen DNA constructs comprising both 5′ regulatory sequences and CDS, originating from both model and non-model organisms. To validate this feature, we sourced data from the literature and evaluated the predicted outcomes via our algorithms.

One such data set we used is from a recent publication^24^, which delved into the role of codon variations within CDSs and how these affect gene expression in *E. coli*. The constructs used in this research shared a fixed promoter and 5′ UTR, but had differing CDSs, generated from a synonymous codon-randomised library using a red fluorescent protein. Although the theoretical variety of such codon-optimised variants is substantial, the authors were able to characterise the fluorescent outputs of 1459 constructs rigorously in *E. coli*. Importantly, this specific dataset had not been seen by our algorithms prior to this analysis. Employing our platform, we predicted the expression levels of the top ten scoring constructs, which are depicted in Figure 7A (Spearman’s ρ = 0.62 p < 0.01).

**Figure 7.**
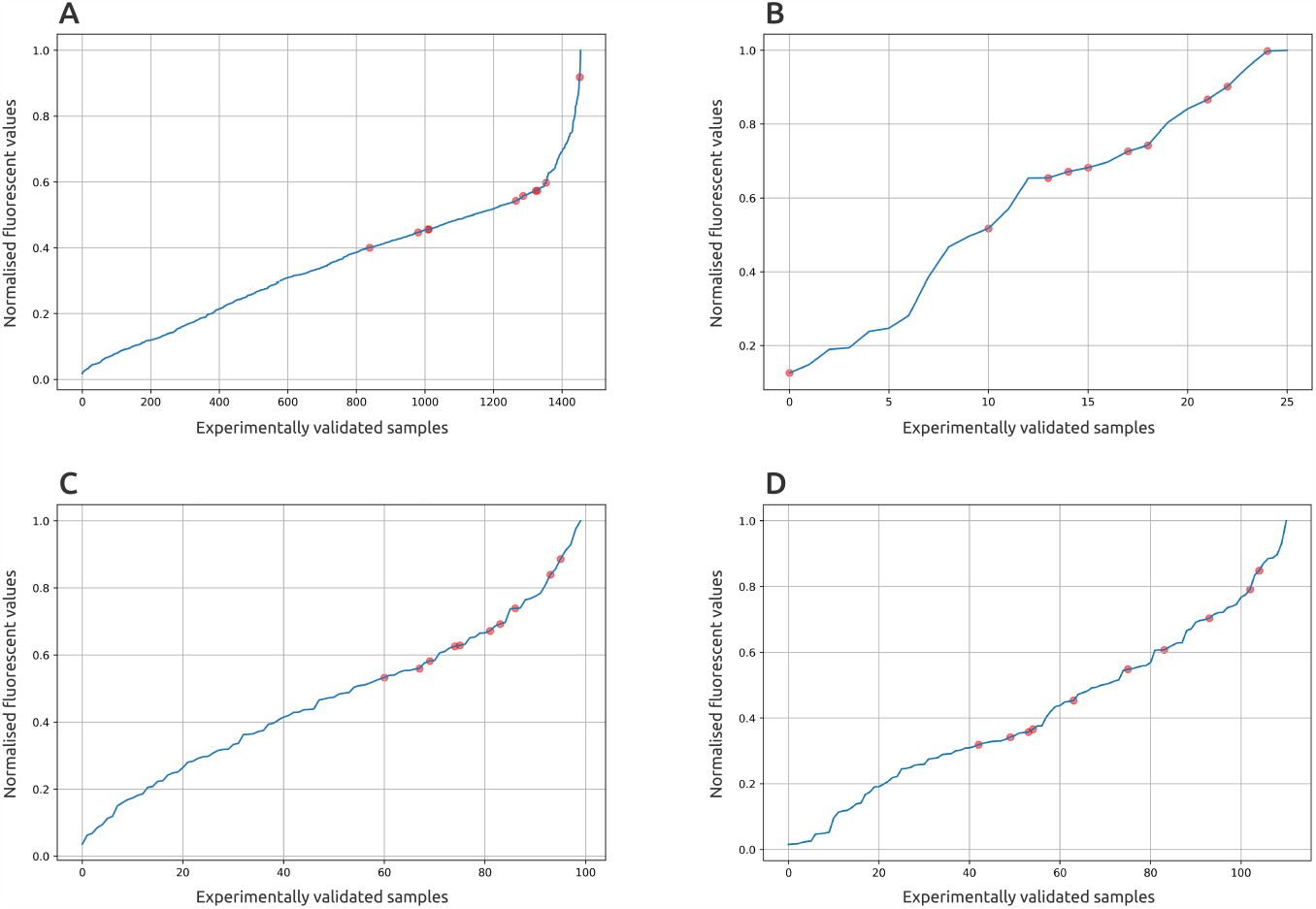
Experimentally validated samples curated from the literature. The top perfmorming 10 constructs based on the predictions performed by the platform are indicated by full red circles. **A** Red fluorescent protein with varying coding sequences characterised in *E. coli*. **B** Promoter and 5′ variants characterised in *V. natriegens* using GFP as the reporter protein. **C** Promoter variants characterised in *S. venezuelae* using GFP as the reporter protein. **D** Promoter variants characterised using GFP in *B. subtilis*. The red spheres indicate the predicted top scoring 10 constructs by the DNA design platform. Spearman’s rank correlation values are **A** 0.62 (p < 0.01), **B** 0.46 (p < 0.01), **C** 0.45 (p < 0.01), **D** 0.40 (p < 0.01) between model predictions and expression values.

To extend beyond *E. coli*, additional datasets from three separate studies have been incorporated. A study by Tietze et al. reports the characterisation of several artifical promoters and 5′ UTRs in *Vibrio natriegens*^27^. This study utilises GFP expression for phenotypic characterisaion of 25 different DNA constructs featuring diverse 5′ regulatory regions. Our platform’s predictive scoring for these constructs is visualised in Figure 7B, and except for two outliers, the constructs that our platform predicted were among the highest performing ones (Spearman’s ρ = 0.46, p < 0.01). The next dataset originates from research focused on constructing a high-yield *Streptomyces venezuelae* in vitro transcription-translation system, and it characterises expression levels of various promoter and 5′ UTR combinations using GFP^28,29^. Our platform also provided phenotype predictions for this data (Figure 7C, Spearman’s ρ = 0.45, p < 0.01).

The final dataset is drawn from a study that investigates the characterisation and application of native phase-dependent promoters in *Bacillus subtilis*^30^. Here, too, our platform made predictions with acceptable accuracy levels (Figure 7D, Spearman’s ρ = 0.40, p < 0.01).

It should be noted that while the predicted expression levels are not exact values, they are estimations that assist in making informed decisions. These results validate the algorithm’s effectiveness in identifying top-performing constructs from a candidate pool. Despite minor shortcomings, the models adeptly capture vital elements necessary for expression level prediction. In summary, the platform demonstrates scalability and efficacy in both de novo design and phenotype prediction across a variety of bacterial hosts.

### Deciphering the Black Box

Employing state-of-the-art AI algorithms, the DNA design platform has the capacity to de novo design of DNA sequences while also predicting their resulting expression levels, thereby offering a comprehensive solution for synthetic biology applications. While the Transformer models display remarkable proficiency in designing DNA sequences and predicting expression levels, their inner workings remain largely inscrutable, necessitating further exploration to fully understand their decision-making mechanisms.

To investigate the model’s weights, we employed Integrated Gradients, an axiomatic attribution method for deep networks, and partially displayed the relevant input sequence sections in Figure 8. Our model suggests that the sequence portions marked in red are suboptimal and result in lower expression scores, whereas those highlighted in green contribute to higher scores. This opens up the possibility of rationally mutating only the red sections to enhance expression levels, rather than introducing mutations over uninformed sections at random. These observations align with a published study where the results were experimentally verified.

**Figure 8.**
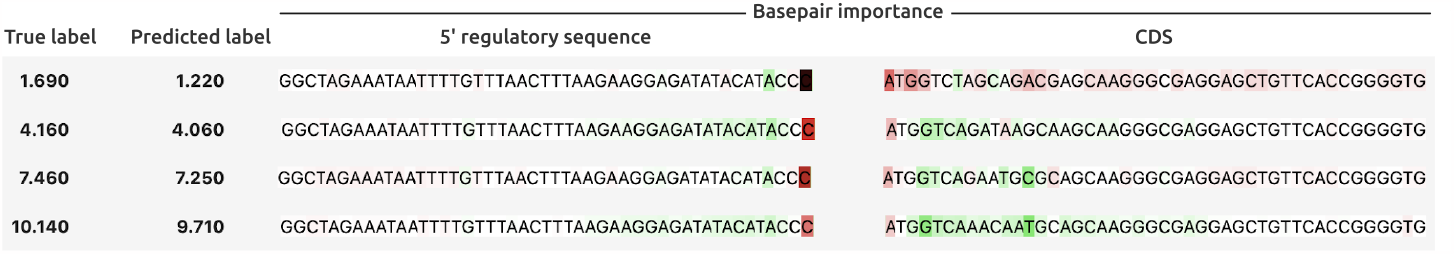
Network analysis of the de novo designed sequence with low score. The green coloured sequences contribute positively, while the red marked sequences lead to lowering the total score. The sequence presented at bottom has the low performing nucleotides mutated hence displayes a higher total score.

This analysis grants us unprecedented insight into the nuanced roles played by specific nucleotides in influencing microbial phenotypes. By unravelling these complexities, we not only enhance the interpretability and trustworthiness of the platform, but also advance our understanding of the nuanced interactions between sequence and function. This pivotal step bridges the gap between machine learning predictions and biological rationale, making the platform an invaluable tool for deciphering the inner workings of our models.

## Discussion and conclusion

The field of biotechnology is experiencing a renaissance, particularly in the post-COVID era, driven by the global push for sustainability and the need to transition towards environmentally friendly biomanufacturing solutions. However, despite the soaring interest the traditional biomanufacturing methods heavily rely on trial-and-error approaches, often neglecting the crucial role of DNA context in achieving desired functionality. This limitation leads to a lack of effective and predictable DNA design, necessitating resource-intensive high-throughput screening efforts. Although proven effective, such approaches are not scalable due to limitations in the high cost of DNA synthesis and the length of the synthesised DNA. To address these challenges and accelerate the field of biomanufacturing, we explore the integration of engineering principles and the predictive power of AI, specifically deep learning transformer models. By incorporating DNA context awareness, the DNA design platform overcomes the limitations of pre-characterised DNA parts, reliance on single codon-optimised CDSs, model organisms, and the absence of predictability. Through DNA sequence analysis and interpretation, we enable informed decision making, optimise genetic element arrangement, and achieve desired outcomes. With predictability, rational DNA design significantly reduces the reliance on trial-and-error, streamlines experimental efforts, and offers a pathway towards scalable and efficient biomanufacturing. Our work highlights the transformative potential of merging biology and AI, ushering in a predictive engineering field and contributing to the development of sustainable and innovative biotechnology solutions.

### Future perspective

While not being the core focus of the reported study, we are compelled to speculate on the foundational principles of scientific investigation. Traditionally, the scientific method has been grounded in rational enquiry, prioritising the understanding of ‘why’ and ‘how’ before delving into the ‘what’. The predictive capabilities of AI are now making us to reconsider this long-standing approach into question. With AI-generated DNA designs, we find ourselves at an important moment where the necessity for a deep-rooted scientific comprehension of DNA composition may be bypassed. Instead, we can place a high level of trust in the AI-derived schematics. This represents not merely a technological milestone, but also a reconsideration of the core tenets of the scientific methodology.

## Materials Methods

### Bacterial strains and growth conditions

*Bacillus subtilis* cells were grown in Nutrient agar (NA) (Oxoid) at 37 °C, supplemented with 50 μg/ml kanamycin. *Corynebacterium glutamicum* cells were grown in Brain Heart Infusion Broth (BHIB, 37 g/l Brain Heart Infusion mix [Difco] and 91 g/l sorbitol) or Brain Heart Infusion Agar (BHIA, 37 g/l Brain Heart Infusion mix [Difco] and 10 g/l agar) at 30°C, supplemented with 15 μg/ml chloramphenicol.

*Escherichia coli* cells were grown in lysogeny broth (LB, 10 g/l tryptone, 5 g/l yeast extract and 5 g/l NaCl) or lysogeny agar (LA, LB + 15 g/l agar) at 37 °C, supplemented with 50 μg/ml kanamycin. For induction of the ChnR/Pb system, cyclohexanone (Sigma) was added at the concentration of 1 mM.

*Pseudomonas putida* cells were grown in LA or LB at 30 °C, supplemented with 50 μg/ml kanamycin.

### Host uniqueness

For this analysis we used the HGTree v2.0 orthosets data to identify orthologs genes. We downloaded data from http://hgtree2.snu.ac.kr/download/orthosets/bacteria/. We restricted the groups to those with exact amino acid sequence in our own datasets. We found >12000 groups based on this and out of those we selected top 35 groups based on the match counts for same amino acids among different hosts for a given group. Out of those we selected 4 group with Uniprot IDs: A0A064BZB4, A0A0E1M7N8, A0A1C1EV68 and A0A0J6J4S5.

### Codon usage

Codon usage analysis is performed by looking at the first 100 codons (300 bp) for highly expressed genes, top 30% expressed genes based on RNA-seq dataset in *E. coli, C. glutamicum, S. venezualae* and *B. subtilis*.

### Codon optimisation

We generate upto 10 000 candidates for each given gene sequence per host and all candidates need to pass constraints defined by us using DNA Chisel^25^. Afterwards we rank them based on the given regulatory sequence for the construct to select the best optimised coding sequences set ca 100.

## Supporting information

Supplementary file with DNA sequences

## Supplementary

**Figure S1.**
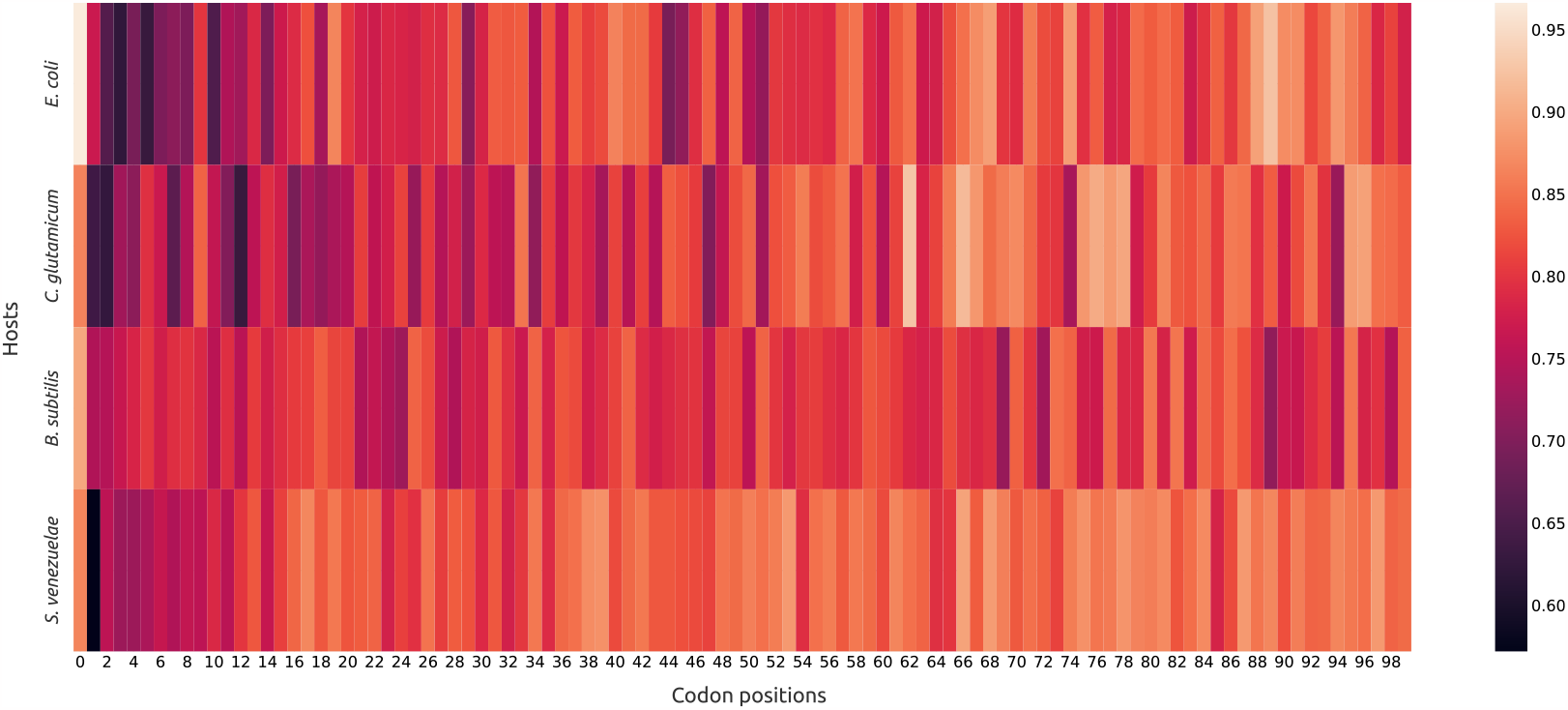
Codon usage analysis of first 100 codons (300 bp) for highly expressed genes, top 30% expressed genes based on RNA-seq dataset in *E. coli, C. glutamicum, S. venezualae* and *B. subtilis*. The color gradient indicates rare codons from dark to abundant codons in light colour.

For the protein ID A0A064BZB4 we have identified 15 different *Streptococcus* species that has the same CDS but different regulatory sequences (Figure S2). The identified 15 hosts are *S. pneumoniae, S. suis, S. oralis, S. mitis, S. parasanguinis, S. sanguinis, S. acidominimus, S. sp*., *S. cristatus, S. australis, S. gordonii, S. ruminantium, S. viridans, S. infantis*, and *S. pseudopneumoniae*.

**Figure S2.**
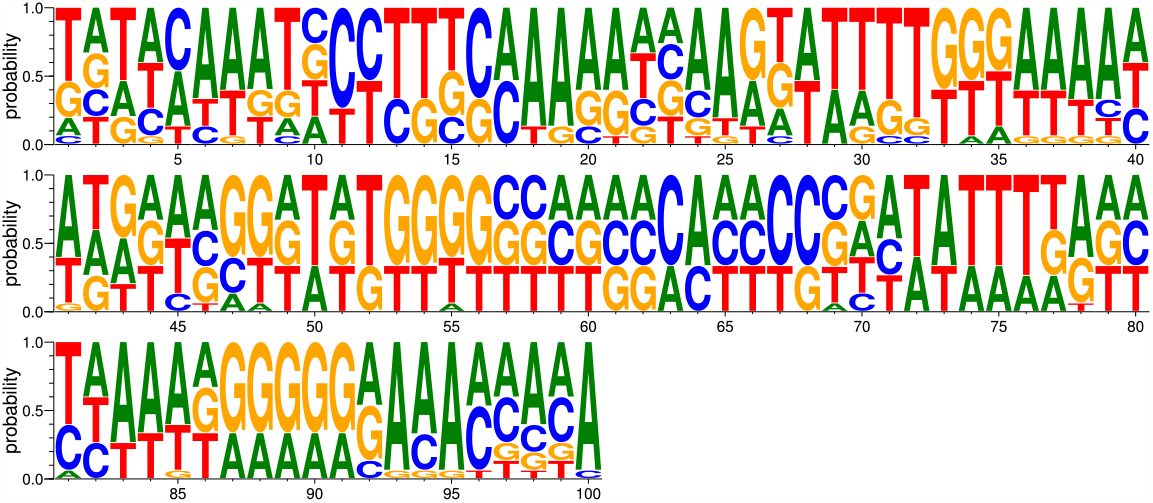
The regulatory sequence of A0A064BZB4 in 15 different *Streptococcus* species.

For the protein ID A0A0E1M7N8 we have identified 15 different *Bacillus* species that has the same CDS but different regulatory sequences (Figure S3). The identified 15 hosts are *B. cereus, B. wiedmannii, B. thuringiensis, B. sp*., *B. anthracis, B. subtilis, B. mobilis, B. thuringiensis*.

**Figure S3.**
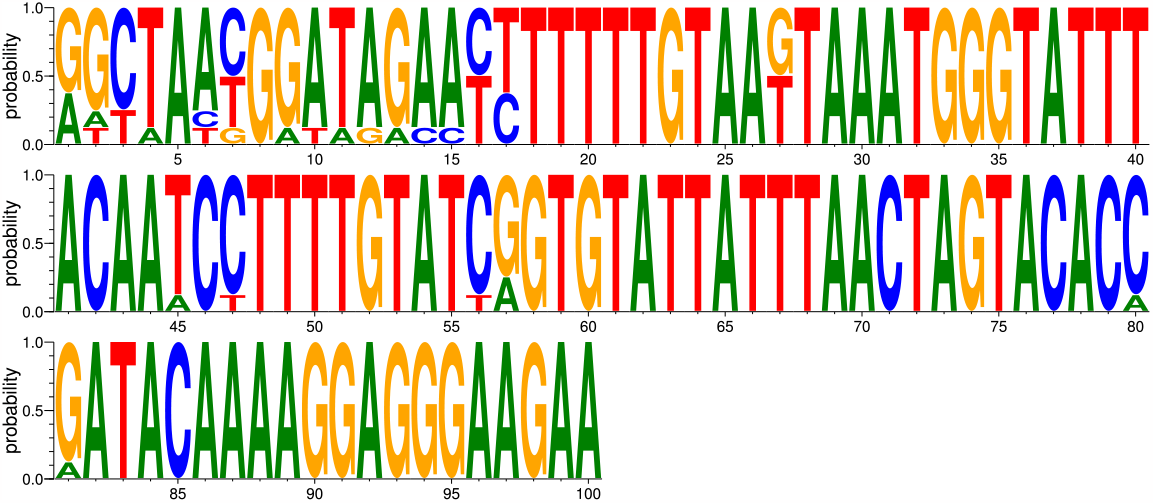
The regulatory sequence of A0A0E1M7N8 in 8 different *Bacillus* species.

For the protein ID A0A0J6J4S5 we have identified 15 different *Pseudomonas* species that has the same CDS but different regulatory sequences (Figure S4). The identified 15 hosts are *Pseudomonas deceptionensis, P. kribbensis, P. sp*., *P. arsenicoxydans, P. versuta, P. baetica, P. fluorescens, P. fragi, P. reinekei, P. mohnii, P. putida, P. mandelii, P. silesiensis, P. prosekii, P. lundensis*.

**Figure S4.**
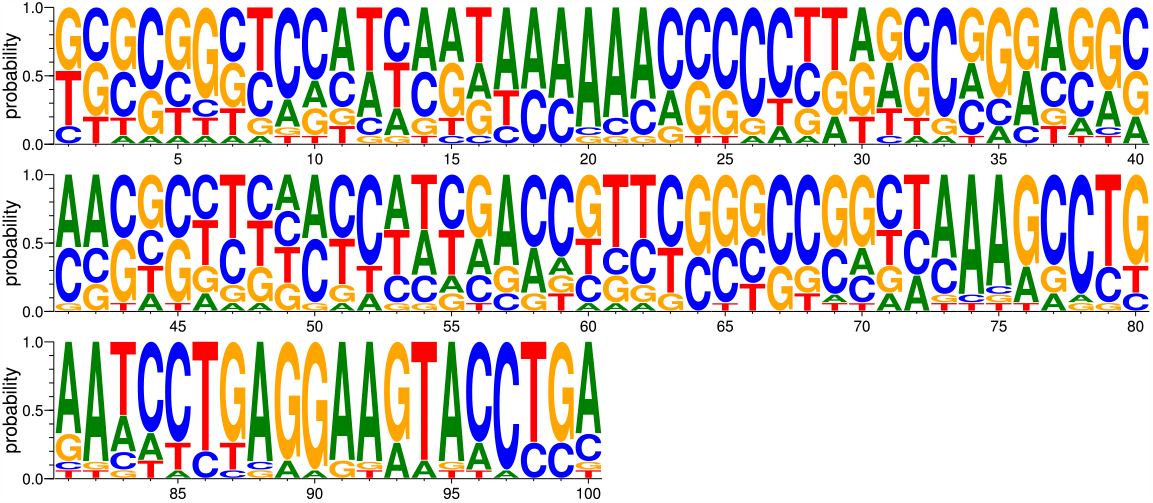
The regulatory sequence of A0A0J6J4S5 in 12 different *Pseudomonas* species.

For the protein ID A0A1C1EV68 we have identified 15 different species that has the same CDS but different regulatory sequences (Figure S5). The identified 15 hosts are *Klebsiella quasipneumoniae, Klebsiella sp*., *K. quasivariicola, K. variicola, K. aerogenes, K. pneumoniae, Enterobacter sp*., *Citrobacter freundii, C. sp*., *C. amalonaticus, C. farmeri, Enterobacteriaceae bacterium*.

**Figure S5.**
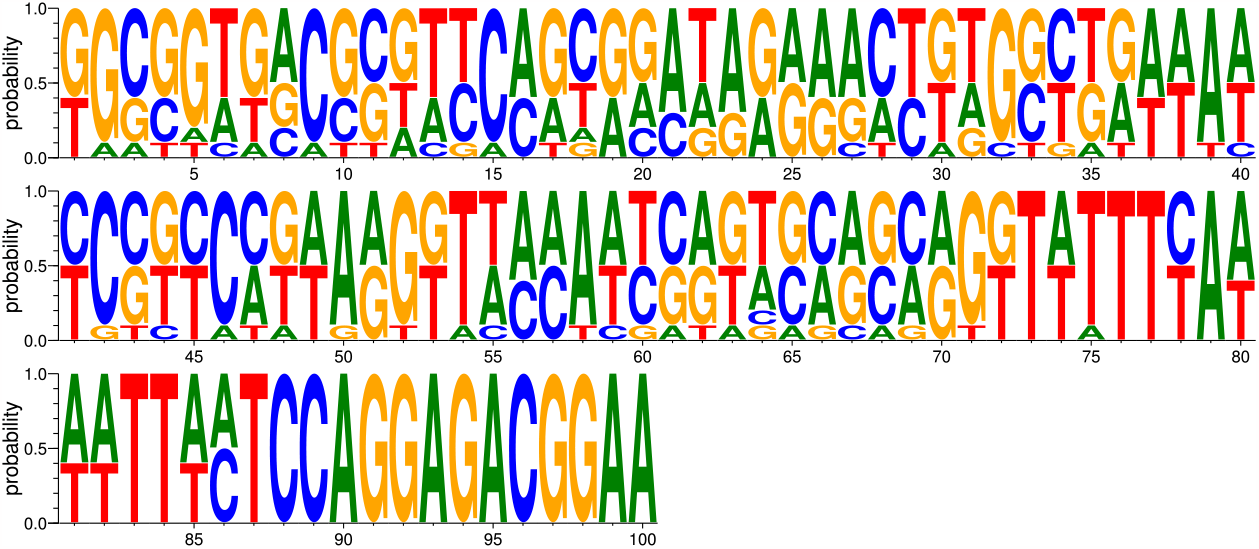
The regulatory sequence of A0A1C1EV68 in 15 different bacterial species.

**Figure S6.**
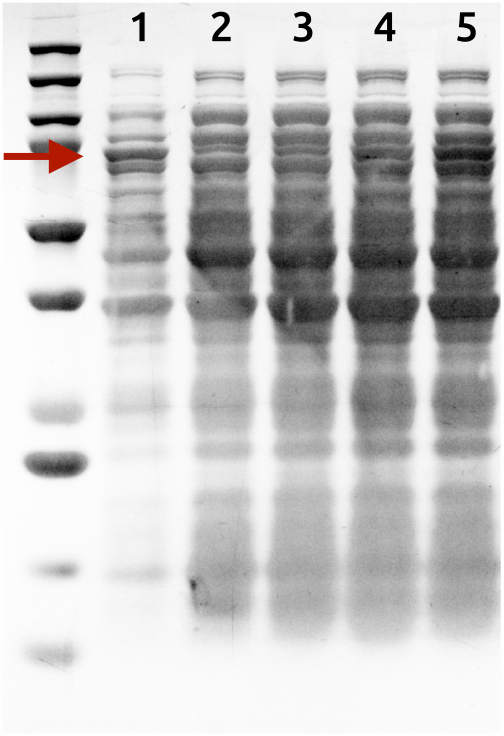
Five DNA constructs with ChnR/Pb systems expressing M-MulvRT.

